# Scorpionfish BPI is highly active against multiple drug-resistant *Pseudomonas aeruginosa* isolates from cystic fibrosis patients

**DOI:** 10.1101/2023.02.19.529133

**Authors:** Jonas M. Holzinger, Martina Toelge, Maren Werner, Katharina U. Ederer, Heiko Siegmund, David Peterhoff, Stefan Blaas, Nicolas Gisch, Christoph Brochhausen-Delius, André Gessner, Sigrid Bülow

## Abstract

Chronic pulmonary infection is a hallmark of cystic fibrosis (CF) and requires continuous antibiotic treatment. In this context, *Pseudomonas aeruginosa* (*Pa*) is of special concern since colonizing strains frequently acquire multiple drug resistance (MDR). Bactericidal/permeability-increasing protein (BPI) is a neutrophil-derived, endogenous protein with high bactericidal potency against Gram-negative bacteria. However, a significant range of CF patients produce anti-neutrophil cytoplasmic antibodies against BPI (BPI-ANCA) thereby neutralizing its bactericidal function. In accordance with literature, we describe that 54.3% of a total of 46 CF patients expressed BPI-ANCA. Importantly, an orthologous protein to human BPI (huBPI) derived from the scorpionfish *Sebastes schlegelii* (scoBPI) completely escaped recognition by these autoantibodies. Moreover, scoBPI exhibited high anti-inflammatory potency towards *Pa* LPS, and was bactericidal against MDR *Pa* derived from CF patients at nanomolar concentrations. In conclusion, our results highlight the potential of highly active orthologous proteins of huBPI in treatment of MDR *Pa* infections, especially in the presence of BPI-ANCA.

## Introduction

Cystic fibrosis (CF) is an autosomal recessive disorder induced by loss-of-function mutations in the gene encoding for cystic fibrosis transmembrane conductance regulator (CFTR) protein (1). Defects in *CFTR* negatively influence homeostasis of airway surface liquid and mucus of the respiratory epithelium, aggravating mucociliary clearance (2,3). Despite the clinical improvements achieved by CFTR modulators in people with at least one F508del mutation (4), chronic pulmonary infections with the opportunistic pathogen *Pseudomonas aeruginosa* (*Pa*) will continue to determine disease progression in untreated or unresponsive CF (5,6). In these cases, continuous suppressive antibiotic therapy is a standard regime to improve survival. However, the long-term application of antibiotics promotes the emergence of persister populations (7,8) and multiple drug-resistant (MDR) bacteria (9), thus limiting the therapeutic options. The neutrophil-derived bactericidal/permeability-increasing protein (BPI) is highly active against Gram-negative bacteria (10,11). Additionally, BPI acts as an anti-inflammatory mediator by neutralizing Gram-negative lipopolysaccharide (LPS; 12,13). By inhibiting growth of *Escherichia coli* (*Ec*) and by neutralizing endotoxic activity of LPS in a picomolar range (14), it is one of the most effective antimicrobials in the human body. The recombinant N-terminal fragment of human BPI (rBPI21) was shown to be highly effective against MDR Gram-negative bacteria including MDR *Pa* (15). However, anti-neutrophil cytoplasmic autoantibodies directed against BPI (BPI-ANCA) found in up to 83% of CF patients (16) inhibit the bactericidal function of BPI (17,18). Coherently, BPI-ANCA correlate with increased colonization with *Pa* (19), impaired lung function and poor prognosis (20). To restore BPI function, we speculated that orthologous proteins with conserved bactericidal and LPS-neutralizing activity would escape recognition by intrinsic BPI-ANCA and therefore preserve their bactericidal activity when applied to BPI-ANCA positive CF patients. Based on known properties, sequence homologies and structural considerations we selected murine BPI (muBPI) of the house mouse *Mus musculus* (21), scorpionfish BPI (scoBPI) of the Korean rockfish *Sebastes schlegelii* (22) and oyster BPI (osBPI) of *Magallana gigas* (Pacific oyster, previously *Crassostrea gigas;* 23,24) for comparison to human BPI (huBPI, *Homo sapiens*). In contrast to huBPI, neither muBPI nor scoBPI were recognized by BPI-ANCA present in the sera of CF patients. Furthermore, scoBPI displayed superior LPS-neutralizing capacity and bactericidal activity towards MDR *Pa* at a nanomolar range.

## Results

### Comparison of the N-terminal barrel in orthologous BPI

The N-terminal domain of huBPI contains three highly cationic regions (region I to III; Fig. 1A, B), which engage with the negatively charged moieties of LPS to neutralize its pro-inflammatory potential (25,26). Therefore, we hypothesized that the affinity towards LPS is roughly determined by the respective quantity of basic amino acids. Preserved LPS neutralization as well as bactericidal activity was previously described for BPI orthologues derived from distantly related species such as the Pacific oyster (23,24) and ray finned fish (Actinopterygii; 22,27). Upon 10 published sequences of Actinopterygii BPI, scoBPI comprised the highest number of cationic arginine, histidine, and lysine amino acids in the corresponding regions (Fig. S1A-C). A sequence alignment also revealed that scoBPI contained more basic amino acids in regions I to III than muBPI and osBPI (Fig. 1A, C, D). In contrast to LPS binding, only 15 amino acids in region II determine the antimicrobial activity in huBPI (25; Fig. 1A). Within this region, sequence analysis revealed a conserved cationic charge with a total of five basic amino acids for muBPI or six for osBPI, scoBPI, huBPI, respectively (Fig. 1A). Exploiting the published three-dimensional structures and electrostatic surface potentials for huBPI (28) and AlphaFold Protein Structure Database (29,30) predictions for the orthologous proteins displayed a highly condensed positively charged cluster in scoBPI that was exclusively found at the N-terminal tip (Fig. 1E). Within the investigated class of Actinopterygii, acBPI of the sterlet *Acipenser ruthenus* (subclass Chondrostei) and gaBPI of the Atlantic cod *Gadus morhua* (subclass Neopterygii) also presented a cluster of positive charges in the respective area along with a relatively high quantity of basic amino acids in functional regions I – III (Fig. S1B, S2A). Interestingly, overall sequence identity with scoBPI was comparably low (56.4% and 62.8%, respectively; Fig. S2B). In conclusion, this initial screen revaled a higher quantity of positively charged surface areas for scoBPI and other fish BPI as compared to the mammalian muBPI or the distantly related osBPI.

**Figure 1:**
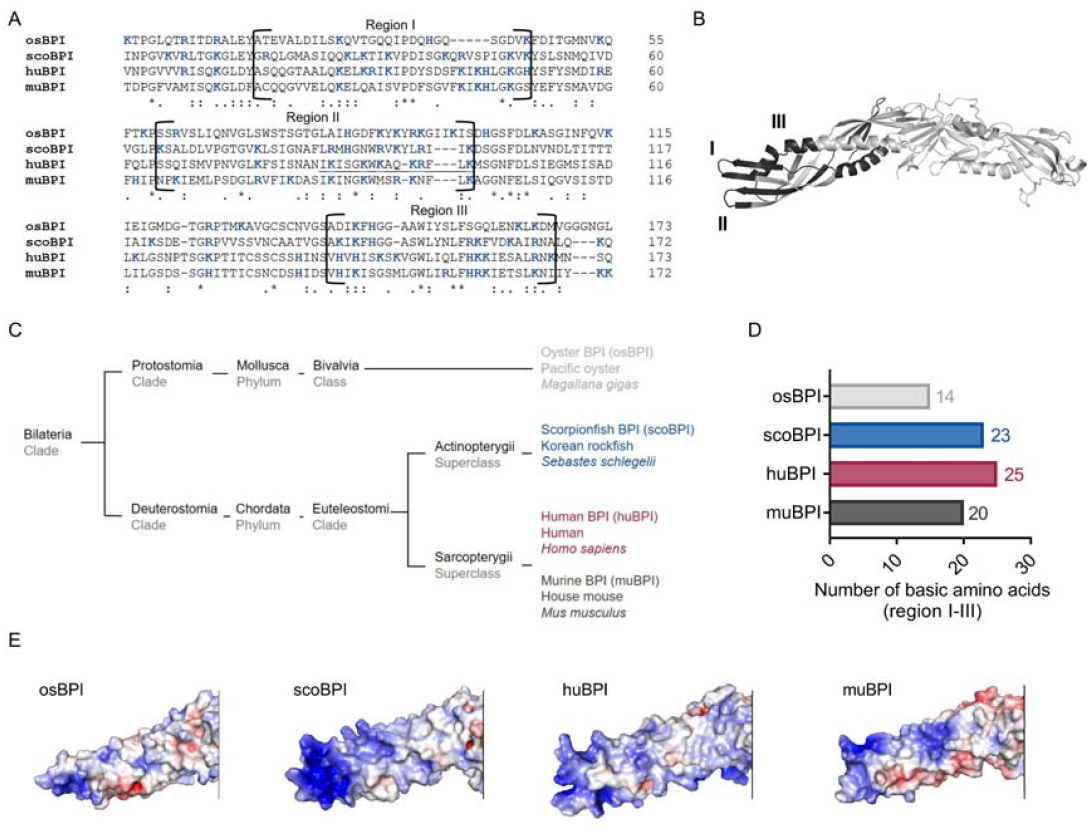
Sequence alignment and analysis of orthologous proteins to huBPI. (**A**) Sequence alignment of amino acid sequences of orthologous proteins. Functional regions I, II and III are framed and positively charged amino acids are shown in blue. The amino acid sequence mediating the bactericidal activity of huBPI is underlined. (**B**) Functional region I, II and III (dark grey) as shown for huBPI. (**C**) Phylogenetic tree analysis mapping BPI orthologous proteins to their respective taxonomic ranks. (**D**) Number of basic amino acids in regions I–III as shown for osBPI, scoBPI, huBPI and scoBPI. (**E**) Electrostatic surface potentials calculated for the N-terminal barrel of osBPI, scoBPI, huBPI and muBPI. Negatively charged areas are colored in red, positively charged domains are shown in blue.

### No recognition of scoBPI by BPI-ANCA from CF patients

As stated, the recognition of endogenous BPI by BPI-ANCA is known to inhibit its bactericidal function and to predict disease progression in CF patients (20). Sequence homology between huBPI and Actinopterygii orthologues, including scoBPI, was low ranging from 36.3% to 38.9% (Fig. S2B), and, consistent with distant or close relationship to huBPI, 24.9% for osBPI and 54.3% for muBPI, respectively (Fig. 2A). Therefore, we speculated that BPI-ANCA in sera of CF patients would not bind to non-huBPI. First, we screened the sera of 46 CF patients for presence of BPI and BPI-ANCA. In accordance with increased levels of BPI in respiratory secretions of CF patients and during the course of respiratory infections (31,32), levels of BPI in sera were significantly elevated (control: mean 75.14 ng/ml, 95% CI 41.4 – 108.9 ng/ml; CF: mean 1115.0 ng/ml, CF 95% CI 0.0 – 2232.0 ng/ml; p < 0.0001;) indicating neutrophil activation in a portion of the CF cohort (Fig. 2B). Furthermore, BPI-ANCA as measured in arbitrary units (AU) were also significantly increased in the CF patients (control: mean 150 AU, 95% confidence interval (CI) 107.5 – 192.4 AU; CF: mean 1511.0 AU, 95% CI 892.3 – 2131.0 AU, p < 0.0001; Fig. 2C). Given previous publications on bactericidal activity (21–24), we selected muBPI, scoBPI and osBPI for functional analysis. After recombinant expression of huBPI, muBPI, scoBPI and osBPI (Fig. 2D), we used beads coupled with the respective BPI orthologues to analyze BPI-ANCA interaction. While huBPI was bound by anti-BPI antibodies in 54.3%, muBPI and scoBPI were not recognized at all and osBPI was only detected by 6.5% of the CF sera (Fig. 2E). Thus, the orthologous proteins muBPI and scoBPI do not contain epitopes detected by intrinsic BPI-ANCA of the tested cohort of CF patients.

**Figure 2:**
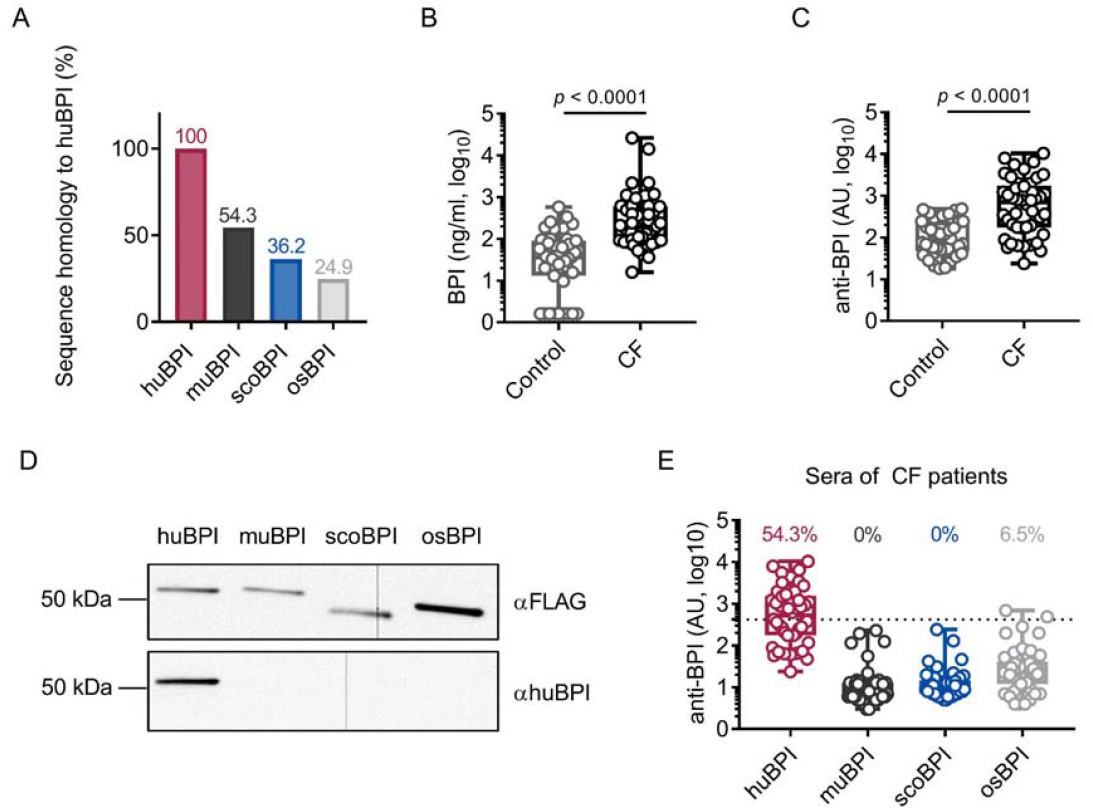
BPI-ANCA from CF patients do not recognize orthologous proteins of huBPI. (**A**) Sequence homology of amino acid sequences to holo huBPI without signal peptide. (**B, C**) Levels of BPI (A) and anti-BPI antibodies indicated in arbitrary units (AU; B) in the sera of 46 individual CF patients and 43 healthy controls. (**D**) Western blot analysis of recombinantly expressed proteins. Bands were visualized with anti-FLAG (αFLAG) or anti-human BPI (αhuBPI) antibodies. One representative experiment of 2 biological replicates using different protein lots in separate assays is shown. (**E**) Recognition of recombinantly expressed proteins by anti-BPI antibodies in the sera of 46 individual CF patients is indicated in AU. The signal cut off is indicated by a dotted line. Data are shown as box plots showing median, upper, and lower quartiles and whiskers indicating minimal and maximal values (B, C, E) or bar plots using calculated values (A). For depiction in the logarithmic scale, values of zero were set to the lower limit of detection of the assay (B). Statistical testing was performed using the Mann-Whitney *U* test. Statistical significance is indicated by *p* values. The online version of this article includes the following source data for Figure 2: **Source data 1**: Raw data for Figure 2B, C and E. Uncropped and labeled blots for Figure 2D.

### Superior anti-inflammatory activity of scoBPI in human PBMCs

Persistent bacterial colonization of CF patients, including Gram-negative bacteria and particularly *Pa*, drives chronic pulmonary inflammation and tissue damage (8). The main immunostimulatory component of Gram-negative bacteria is LPS. Importantly, *Ec*-derived LPS can effectively be neutralized by picomolar concentrations of huBPI (14). Therefore, we wanted to investigate whether the recombinant BPI orthologues were competent to interact with *Ec* LPS and neutralize its immunostimulatory potential. Importantly, we found that secretion of IL-6 in human PBMCs by *Ec* LPS was most effectively inhibited by scoBPI, exceeding huBPI, muBPI and osBPI (Fig. 3A). This superior LPS-neutralizing capacity compared to huBPI could be observed for low picomolar concentrations of scoBPI (Fig. 3B). To explore interaction of huBPI or orthologues with LPS derived from *Pa*, we purified LPS of *Pa* PAO1. Similar to incubation of huBPI and LPS derived from *Salmonella minnesota* Re595 (33), incubation of both huBPI and scoBPI with *Pa* LPS fostered prominent increase in aggregate size as measured by DLS (Fig. 3C) indicating close interaction. Congruent with aggregate formation, the pro-inflammatory potential of *Pa* LPS was drastically inhibited by huBPI and scoBPI (Fig. 3D, E). In comparison to huBPI, scoBPI exhibited a superior anti-inflammatory potential as indicated by diminished production of IL-6 and TNF in response to *Pa* LPS. These data suggest that scoBPI holds strong anti-inflammatory properties towards *Ec* and *Pa* LPS.

**Figure 3:**
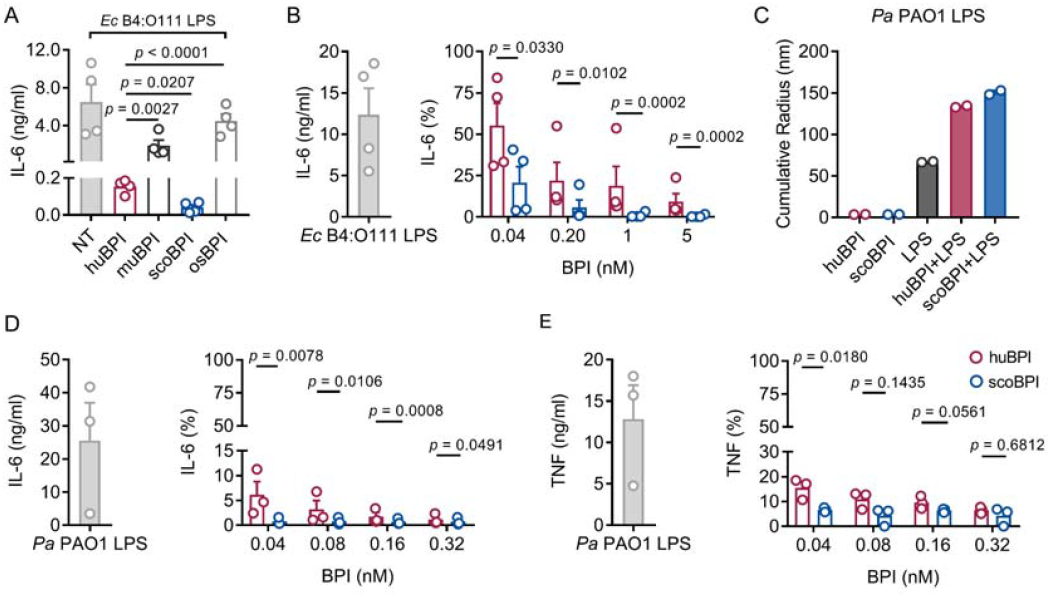
Potent anti-inflammatory action of scoBPI in human and murine immune cells. (**A**) Levels of IL-6 in supernatants of PBMCs stimulated for 24 h with *Ec* B4:O111 LPS (10 ng/ml) ± huBPI, muBPI, scoBPI or osBPI (25 nM; n = 4). (**B**) Levels of IL-6 in supernatants of PBMCs stimulated for 24 h with *Ec* B4:O111 LPS (10 ng/ml) ± huBPI (red) or scoBPI (blue) in concentration as indicated. (**C**) Aggregate size of *Pa* PAO1 LPS ± huBPI or scoBPI as determined by NanoDLS (n = 2). (**D, E**) Quantification of IL-6 (D) and TNF (E) levels in supernatants of PBMCs stimulated for 24 h with *Pa* PAO1 LPS (100 ng/ml) ± huBPI (red) or scoBPI (blue) in concentrations as indicated (n = 3). Experiments were performed using PBMCs of four (A, B) or three (D, E) individual blood donors. NanoDLS experiments were performed as technical replicates in 2 separate assays (C). Data are shown as means ± SEM. Statistical testing was performed using the student’s ratio paired *t* test (A, B, D, E). Statistical significance is indicated by *p* values. The online version of this article includes the following source data for Figure 3: **Source data 1**: Raw data for Figure 3A - E.

### Bactericidal activity of scoBPI against multiple drug-resistant isolates of *Pseudomonas aeruginosa*

To evaluate the antimicrobial activity of the recombinantly expressed orthologous BPI proteins we first compared their capacity to inhibit growth of *Ec* DH10B. Activity of huBPI (IC_50_ 0.032 ± 0.011 nM), scoBPI (IC_50_ 0.032 ± 0.006 nM) and muBPI (IC_50_ 0.061 ± 0.016 nM) were in low picomolar range, while osBPI (IC_50_ 60.16 ± 21.43 nM) required higher nanomolar concentrations to fully inhibit bacterial growth (Fig. 4A). Bactericidal assays with *Pa* PAO1 revealed impaired activity of muBPI and osBPI, while huBPI and scoBPI potently inhibited bacterial growth (Fig. 4B). Next, we were interested if BPI was also active against the highly adapted *Pa* strains from CF patients. Therefore, six distinct MDR *Pa* isolates of five patients were selected (mean age at sample collection 24.8 ± 7.1 years). Broad-spectrum resistance including resistance to Ciprofloxacin, Piperacillin, Piperacillin/Tazobactam, Ceftazidim, Cefepim, Imipenem, and Meropenem was verified by disc diffusion testing (Table 1). Despite major adaptions during chronic *Pa* infections (8), three of the six isolates were also highly susceptible to both huBPI and scoBPI at concentrations of 20 nM after one hour of incubation (Fig. 4C). More than 50% growth reduction was achieved in all isolates within one hour at a concentration of 500 nM of scoBPI. Thereby, a trend towards superior antibiotic activity against MDR *Pa* for scoBPI compared to huBPI was observed (*p* = 0.0756). One of the less sensitive strains (isolate 5) possibly emerged from a previous isolate (isolate 2) in a time course of 3.3 years, possibly indicating adaptions to a long-term BPI exposure *in vivo*. In accordance with literature for huBPI (31), mucoid strains were susceptible to both huBPI and scoBPI. The time-dependent bactericidal action of scoBPI against the MDR *Pa* isolates was also visualized in bactericidal viability assays (Fig. 4D). Corresponding transmission electron microscopy (TEM) of a representative scoBPI-treated MDR *Pa* isolate revealed severe damage to the bacterial cell wall and efflux of cytoplasm in affected bacteria (Fig. 4E, Fig. S3). In conclusion, scoBPI is bactericidal towards MDR *Pa* isolates derived from chronically infected CF patients.

**Figure 4:**
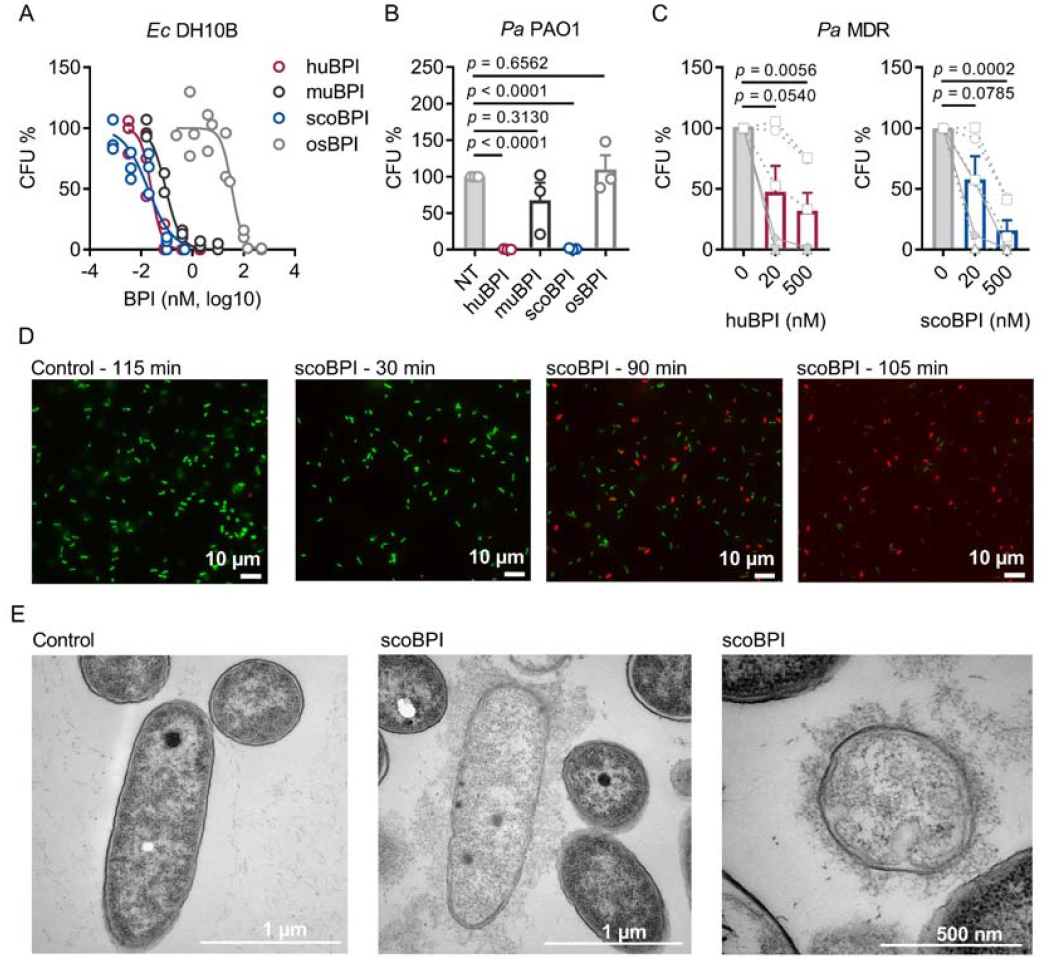
Bactericidal activity of huBPI and scoBPI against *Ec* and MDR isolates of *Pa*. **(A)** Dose-response curves for *Ec* DH10B incubated with increasing concentrations of huBPI, muBPI, scoBPI or osBPI (n = 3). **(B)** Bactericidal activity of huBPI, muBPI, scoBPI and osBPI 500 nM against *Pa* PAO1 (n = 3). **(C)** Antibacterial activity of huBPI and scoBPI at concentrations of 20 and 500 nM against six MDR isolates of *Pa* obtained from five individual CF patients. Isolates (isolate 2 and 5 in Table 1) that originate from the same patient are shown as a square. Mucoid isolates are displayed as filled symbols connected by continuous lines. (**D**) Images of a bacterial viability assay performed with one representative *Pa* MDR strain. Viable bacteria are seen in green and dead bacteria in red, respectively. Bar scales for 10 μm. **(E)** Electron transmission microscopy of the representative *Pa* MDR isolate shown in (D) after treatment with PBS (control) or scoBPI. Bar scales for 1 μm (left, middle) or 500 nm (right). Data are shown as individual values (A) or means ± SEM (B, C) of technical replicates using the same lots of BPI and bacteria in 3 separate assays (A, B). Statistical testing was performed using the student’s paired *t* test (A-C). Statistical significance is indicated by *p* values. The online version of this article includes the following source data for Figure 4: **Source data 1**: Raw data for Figure 4A - C.

**Table 1:**
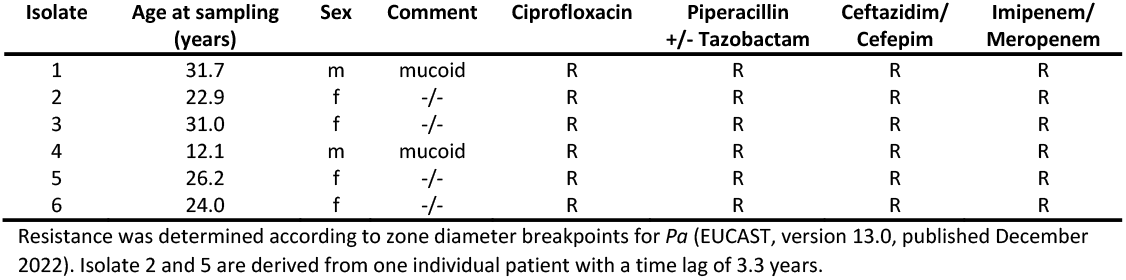
Characteristics of CF patients and MDR *Pa* isolates.

## Discussion

Here we demonstrate that scoBPI, an orthologous protein to huBPI, escapes recognition by BPI-ANCA of CF patients, exhibits high anti-inflammatory potency towards LPS and exhibits bactericidal activity against *Pa*, including MDR *Pa* derived from CF patients. Indeed, we even observed a trend towards superior antibiotic activity against MDR *Pa* for scoBPI compared to huBPI at a concentration of 500 nM. As opposed to established antibiotics that mainly target processes associated with bacterial cell envelope biogenesis, DNA replication, transcription, and protein biosynthesis, the mode of action of BPI relies on membrane disruption by binding to LPS of the outer cell wall in Gram-negative bacteria (34). Bactericidal activity via membrane perturbation of MDR *Pa* by scoBPI could be verified by TEM and explains the susceptibility to BPI despite MDR. As seen by viability imaging, the bactericidal activity of scoBPI was impressively fast and occurred within one to three hours. Activity via membrane permebilisation implies broad inhibition ranging from non-replicating bacteria, such as persister mutants of *Pa* commonly found in CF patients (35), to fast growing *Pa* populations which avoid antibiotic accumulation (36). Sustained immune responses towards *Pa* in CF patients trigger adaption of a mucoid phenotype that is associated with resistance to host antimicrobials (37), impaired phagocytosis (38) and formation of biofilms (39), allowing for bacterial persistence in the CF environment. As shown previously for huBPI (31), mucoid *Pa* isolates were indeed highly susceptible to the bactericidal action of scoBPI. Beyond CF, ESKAPE pathogens (*<u>E</u>nterococcus faecium*, *<u>S</u>taphylococcus aureus*, *<u>K</u>lebsiella pneumoniae*, *<u>A</u>cinetobacter baumannii*, *<u>P</u>a* and *<u>E</u>nterobacter* spp.) were defined by the WHO as priority pathogens for new antimicrobial drug development as these are continuously associated with multiple drug resistance and severe nosocomial infections (40). In this aspect, further studies of scoBPI on the Gram-negative members of the ESKAPE cluster, such as *Pa*, *Acinetobacter baumanii*-complex, and various Enterobacteriaceae, are worthwhile. It is noteworthy that BPI-ANCA are also frequently found in other inflammatory disorders like non-CF bronchiectasis (16), vasculitis (41) and systemic lupus erythematosus (SLE; 42). Since pulmonary *Pa* infections are common in non-CF bronchiectasis (16) and Gram-negative infections are known to be a leading cause of hospitalization in SLE patients (43), scoBPI might be of interest in patient groups other than CF as well. Administration of the recombinant human rBPI23 (amino acid 1-199) and rBPI21 (amino acid 1-193) have previously been tested in clinical phase I – III studies (44,45). Thereby, rBPI23 efficiently neutralized the endotoxic activity of LPS in healthy volunteers (45). Despite the peracute disease course of pediatric meningococcal sepsis and a limited number of patients, rBPI21 was shown to improve clinical outcome by trend (44). In a less acute setting, such as chronic persisting or exacerbation of *Pa* infections, the use of rBPI21, scoBPI or the N-terminal barrel of scoBPI might even be more promising. Despite long-term infections in the tested MDR *Pa* and varying sensitivities, complete resistance to BPI was not found. Clearly, further studies should address this issue in a larger cohort. As suggested by our PBMC data, scoBPI potently neutralized the immunogenic potential of *Pa*-derived LPS and consequently prevented LPS-induced release of TNF and IL-6. Chronic pulmonary inflammation arises early in CF patients, leading to tissue destruction, and eventually accounts to respiratory failure and the premature deaths of patients (46). The abundance of neutrophil-derived elastase in the lungs of CF patients is considered as the essential mediator of this persistent local inflammation (47) and there is evidence that elastase release is primed by TNF (48). It is also known that macrophages in CF patients are hyperresponsive to LPS, further driving dysregulated inflammatory responses (49). Therefore, the benefits of administration of scoBPI by joining antimicrobial activity and LPS neutralization, including reduction of TNF release, could potentially extenuate inflammation and, thereby, tissue destruction. While LPS binding and the antimicrobial activity of BPI are mediated by the N-terminal barrel of the protein (25), the C-terminal barrel is known to enhance phagocytosis by opsonization (50). Fittingly, murine *Bpi*-deficient neutrophils display decreased phagocytosis of *Pa* in pulmonary infection models which could be reconstituted by exogenous BPI (51). Future *in vitro* studies will elucidate if the C-terminal barrel of scoBPI also opsonizes *Pa* for phagocytosis by human cells. Concerning clinical use, one limitation might be a possible induction of antibodies against scoBPI during long-term or repeated applications. Therefore, therapeutic use may rather be indicated in short-term settings. Moreover, the starlet *Acipenser ruthenus* (Actinopterygii, subclass Chondrostei) is a only distantly related to the scorpionfish *Sebastes schlegelii* (Actinopterygii, subclass Neopterygii). Congruently, acBPI shows only low sequence identitiy to scoBPI, thus low epitope overlap. Surprisingly, we found a similar positively charged cluster at the N-terminal tip of both orthologues. Therefore, Actinopterygii BPI, such as acBPI, may also bear therapeutic potentials and should be tested functionally.

Collectively, we define scoBPI as a distantly related, orthologous protein to huBPI that combines LPS-neutralizing capacity and bactericidal activity towards MDR *Pa* isolates. Further investigations on scoBPI and other BPI orthologues are highly encouraged to develop novel therapeutics against Gram-negative MDR infections.

## Methods

### Expression and purification of recombinant BPI orthologues

Templates for recombinant BPI were designed by flanking the corresponding BPI sequences without native signal-peptides (huBPI: amino acids 32 – 487, muBPI: amino acids 28 – 483, scoBPI: amino acids 19 – 473, osBPI: amino acids 20 – 477) with an N-terminal HA-signal peptide and a C-terminal FLAG-tag. Constructs were ligated into a pcDNA 3 (huBPI, muBPI) or pcDNA 3.1(+) (osBPI, scoBPI; Cat# V79020; Thermo Fisher Scientific, Waltham, MA, USA) vector backbone. BPI orthologues were generated as described previously with slight modifications (14,52). Expi293F cells (Cat# A14527, RRID:CVCL_D615; Thermo Fisher Scientific, Waltham, MA, USA) were transfected using the ExpiFectamine 293 Transfection Kit (Thermo Fisher Scientific, Waltham, MA, USA). Recombinant huBPI, muBPI and scoBPI were purified by cation exchange chromatography on a HiTrap SP HP column (Cytiva, Marlborough, MA, USA) followed by size-exclusion chromatography (Superdex 200 increase 10/300 GL column; Cytiva, Marlborough, MA, USA). Due to insufficient binding to the cation exchange column, osBPI was purified by affinity chromatography on an anti-FLAG (Sigma Aldrich, Taufkirchen, Germany) coupled NHS-activated HP column (Cytiva, Marlborough, MA, USA) followed by size exclusion chromatography.

### Western blotting

For western blotting, 12 μl of purified recombinant orthologous proteins at a concentration of 250 nM were loaded onto a TGX FastCast 12% stain-free gel (Bio-Rad, Feldkirchen, Germany). Electrophoresis was conducted at 220 V for 30 min. Proteins were blotted onto a nitrocellulose membrane using the Trans-Blot Turbo TRA Transfer Kit (Bio-Rad, Feldkirchen, Germany). Following protein transfer, membranes were blocked in 5% non-fat dried milk dissolved in Tris-buffered saline with 0.05% Tween20 (TBS-T) for 1 h. Anti-FLAG M2 antibody (Cat# F3165, RRID: AB_259529; Sigma-Aldrich, Taufkirchen, Germany) or polyclonal anti-human BPI antibody (Cat# HM2170, RRID: AB_532911; Hycult Biotech, Uden, Netherlands) were incubated overnight at 4 °C. After three washes with TBS-T, the corresponding secondary, peroxidase-labeled antibodies rabbit anti-mouse (Cat# 711-035-152, RRID: AB_10015282; Dianova, Hamburg, Germany) and donkey anti-rabbit (Cat# 315-035-048, RRID: AB_2340069; Dianova, Hamburg, Germany) were incubated for 45 min at room temperature with three consequent washing steps. Membranes were then incubated with Clarity Western ECL substrate (Bio-Rad, Feldkirchen, Germany) for five minutes. Chemiluminescence was detected with the digital imaging system Chemi Lux Imager (Intas, Göttingen, Germany).

### Bacteria and antimicrobial resistance testing

*Ec* DH10B (Cat# EC0113; Thermo Fisher Scientific, Waltham, MA, USA) and *Pa* PAO1 (ATCC15692, Cat# DSM 22644; DSMZ, Braunschweig, Germany) were cultivated on Columbia blood agar plates (Thermo Fisher Scientific, Waltham, MA, USA). Antimicrobial resistance of MDR *Pa* strains was determined according to zone diameter breakpoints for *Pa* (EUCAST, version 13.0, published December 2022) in the diagnostic laboratory of the Institute of Clinical Microbiology and Hygiene Regensburg (University Hospital Regensburg, Germany). Microbial identification of all strains was confirmed by Matrix-Assisted Laser Desorption/Ionization Time-Of-Flight Mass Spectrometry (MALDI-TOF MS) using the MALDI Biotyper system (Bruker Corporation, Billerica, MA, USA).

### Preparation of LPS

Isolation and purification of *Pa* PAO1 LPS was performed following previously described procedures (53). In detail, bacteria were grown aerobically with shaking (180 rpm) at 37 °C in LB-Broth Base medium (Thermo Fisher Scientific, Waltham, MA, USA) supplemented with 5 g/l of NaCl until an absorbance of approximately 1.1 at 600 nm was reached. Phenol (90%) was added to reach a final concentration of 1% and the resulting suspension was shaken (90 rpm) for 1 h at 37 °C and then at 4 °C overnight. Cells were collected by centrifugation (9,000 g, 20 min, 4 °C) and subsequently washed three times with water (centrifugation conditions as above). The lyophilized pellet (recovery, 2.78 g) was washed with ethanol, acetone (twice), and diethyl ether, and then dried. The pellet was resuspended in water (approximately 14 mg/ml), sequentially treated for 24 h each at RT with DNase/RNase and proteinase K (100 μl of 10 mg/ml solutions per gram dry weight for each enzyme), then underwent dialysis (14-kDa cutoff) and lyophilization (yielding in 610 mg bacterial mass). For hot phenol-water extraction (54), bacteria were resuspended in 45% aqueous phenol (40 ml per g bacteria) and stirred for 30 min at 68 °C. After centrifugation (5,600 × g) for 20 min at 4 °C, the upper water phase was collected. The extraction was repeated with the same volume of water as had been collected. Combined water phases and the phenolic phase were dialyzed against water at RT (14-kDa cutoff). Prior to lyophilization, the dialyzed phenolic phase (PP) was centrifuged (600 × g for 5 min at 20 °C) and divided into supernatant and sediment. The main part of the LPS was recovered from the water phase (184 mg). A portion (20.9 mg) of this LPS preparation was further purified by gel permeation chromatography on Sephacryl S-400 HR (GE Healthcare, Uppsala, Sweden) on a column (2.5 × 120 cm) using a 50 mM ammonium bicarbonate buffer as eluent (55), yielding 6.8 mg. Such material was used in the described experiments.

### Dynamic light scattering

BPI-LPS aggregate size was determined by dynamic light scattering (DLS). Stock solutions of LPS derived from *Pa* PAO1 (500 μg/ml), huBPI and scoBPI (5 μM each) were pre-incubated alone or in the indicated combinations for 30 min at room temperature before filling the capillaries. Light scattering measurements were performed at 20 °C using Prometheus Panta (NanoTemper Technologies GmbH, Munich, Germany).

### Stimulation of human PBMCs

Isolation of human PBMCs was performed as described previously (14). In detail, blood from healthy volunteers was collected in heparinized tubes (Li-Heparin-Gel-Monovette, Sarstedt, Nümbrecht, Germany). The blood was centrifuged in Leucosep tubes containing Ficoll-Paque PLUS (Cytiva Europe GmbH, Freiburg, Germany) at 1,000 × g for 10 min. Leukocytes were collected from the interphase and subsequently washed twice with RPMI medium 1640 (Thermo Fisher Scientific, Waltham, MA, USA). The PBMC pellet was then resuspended in AIM V (Thermo Fisher Scientific, Waltham, MA, USA) and 10^5^ PBMCs were seeded into 96 well plates (Sarstedt, Nümbrecht, Germany) and rested for 3 h prior to stimulation. Stimulants were diluted in AIM V, combined as indicated and pre-incubated in protein LoBind tubes (Eppendorf, Hamburg, Germany) for 30 min at 37 °C. PBMC supernatants were collected after 24 h and stored at −20 °C for cytokine analysis.

### Quantification of cytokines, BPI and BPI-ANCA

Levels of human interleukin (IL)-6 and tumor necrosis factor (TNF) were quantified by Luminex technology (Austin, TX, USA) as described previously (14). Capture and detection antibodies for human IL-6 were from the OptEIA set for human IL-6 (Cat# 55220, RRID: AB_2869045; BD Biosciences, Heidelberg, Germany) and capture and detection antibodies for human TNF were from the OptEIA set for human TNF (Cat# 555212, RRID: AB_2869042; BD Biosciences, Heidelberg, Germany). Signals were detected by Streptavidin-PE (Agilent, Palo Alto, CA, USA) and cytokine concentrations were quantified with the Human Cytokine 16-Plex Protein Standard (Thermo Fisher Scientific, Waltham, MA, USA). For quantification of BPI in human sera, Luminex-beads coupled with anti-BPI antibody 3F9 (Hycult Biotech, Uden, Netherlands) were incubated with patient or control sera over night at 4 °C. BPI was detected with biotinylated anti-BPI antibody 4H5 (Cat# HM2042, RRID: AB_532909; Hycult Biotech, Uden, Netherlands) and Streptavidin-PE (Agilent, Palo Alto, CA, USA). Concentrations were quantified with BPI as a standard extracted from neutrophils purchased from Wieslab AB (Malmö, Sweden). For measurement of BPI-ANCA recombinant proteins were coupled to beads and binding of autoantibodies was detected by a polyclonal PE-labeled anti-human IgG, Fcγ fragment specific antibody (Cat# 109-116-098, RRID: AB_2337678; Dianova, Hamburg, Germany). The signal cut off for positivity was determined using sera of 43 healthy controls and defined as two standard deviations above the mean signal. Autoantibody levels are indicated as AU. Cytokine and autoantibody levels were measured using the Luminex 100 system (Austin, TX, USA) and subsequently analyzed by using LiquiChip Analyzer Software (Qiagen, Hilden, Germany).

### Dose-response experiments

Dose-response experiments were conducted as described previously (14). In brief, bacteria were grown in LB medium at 37 °C and 220 rpm to an optical density (OD_600_) of 0.4. Bacteria were further diluted in PBS (Sigma Aldrich, Taufkirchen, Germany) with 0.01% Tween20 (PBS-T) and incubated with recombinant BPI or PBS-T as mock control for one hour at 37 °C. Dilutions of bacteria were then plated onto blood agar plates and incubated at 37 °C overnight. Colony forming units were enumerated the next day. For MDR isolates of *Pa*, bacteria were incubated with 20 nM and 500 nM BPI, respectively.

### Bacterial viability assay

For bacterial viability assays the LIVE/DEAD *Bac*Light Bacterial Viability Kit from Thermo Fisher Scientific (Waltham, MA, USA) was used. *Pa* isolates were grown to an OD_600_ of 0.2 – 0.8, pelleted and resuspended in PBS to a final OD_600_ of 1.0. Next, 50 μl of cell suspension was diluted with 10 μl of scoBPI to a final scoBPI concentration of 500 nM. For excitation of dyes and microscopic imaging the Keyence All-in-one Fluorescence Microscope BZ-9000 (Keyence, IL, USA) was used.

### Transmission electron microscopy

Bacteria were grown to an OD600 of 0.4. Bacteria were pelletized and resuspended in PBS-T to an OD_600_ of 10. Then, 5 x 10^9^ bacteria were incubated with 2 μM scoBPI for two hours. Bacteria were pelletized, the supernatant replaced by cacodylate-buffered 2.5% glutaraldehyde solution (all reagents from EMS, Science Services, Munich, Germany) and the samples were fixed for 12 h at room temperature. Alginate (Epredia Cytoblock Replacement Reagents, Thermo Fisher Scientific, Waltham, MA, USA) was used to enhance pellet stability for the untreated control. After that, bacteria treated with scoBPI and untreated control were enclosed with 4% low melting agarose. The Lynx microscopy tissue processor (Reichert-Jung, Wetzlar, Germany) was used for the embedding process, including a secondary fixation with 1% osmium tetroxide solution (EMS, Science Services, Munich, Germany), dehydration and infiltration with EPON-mixture (Embed, DDSA, NMA, DP30). The EPON-infiltrated pellets were transfered in silicon moulds and overlaid with fluid EPON-mixture and harden at 60°C in the heating oven. Semithin sections (0,75 μm) were used to define relevant areas and ultrathin sections (80 nm) were then cut accordingly using the Reichert Ultracut S (LEICA Microsystems, Wetzlar, Germany). The ultrathin sections were then contrasted with aqueous 2% uranyl-acetate (EMS, Science Services, Munich, Germany) and 2% lead-citrate solution (Leica Ultrostain II, Leica Microsystems, Wetzlar, Germany) for 10 min each. Electron-microscopic analysis was performed using the LEO 912 AB electron microscope (Zeiss, Oberkochen, Germany).

### Sequence analysis, alignment, and homology calculation

Analyzed protein sequences were from UniProt (56) (huBPI: P17213, muBPI: Q67E05, scoBPI: A0A481NSZ4, osBPI: C4NY84). Sequences were aligned using Clustal Omega (57). Sequence identities were calculated with SIM-alignment tool for protein sequences (58).

### Structure modeling, graphical depictions and statistical analysis

The Protein Data Bank (PDB) structure for huBPI (PDB DOI: 10.2210/pdb1BP1/pdb) deposited by Beamer *et al*. (28) and AlphaFold Protein Structure Database (29,30) predictions for muBPI, scoBPI and osBPI were used for three-dimensional modeling. Three dimensional structures were rendered in PyMOL (PyMOL Molecular Graphics System, Version 2.3.2 Schrodinger, LLC., New York, NY, USA) and electrostatics were calculated using the APBS plugin for PyMOL. Depictions of graphs and statistical analyses were done using GraphPad Prism, version 7.01 (GraphPad Software, San Diego, CA, USA). Results are depicted as indicated in the figure legends as absolute values, means ± SEM or box plots showing median, upper, and lower quartiles and whiskers indicating minimal and maximal values. Statistical tests were performed as described in the figure legends. Thereby *t* test was chosen when values were normally distributed. *P* values < 0.05 were considered statistically significant. Values < 0 of the lower 95% CI were set to 0.

## Ethics Statement

This study was carried out in accordance with the recommendations of the Declaration of Helsinki. Diagnostic leftover samples, stored at the Institute of Clinical Microbiology and Hygiene, University Hospital Regensburg, were used for serum analysis. PBMCs were purified from blood of healthy donors. All donors of PBMCs gave written informed consent. The protocols were approved by the local ethics committee (Ethikkommision an der Universität Regensburg, 18-1269-101 and 16-302_1-101).

## Acknowledgments

We gratefully acknowledge Sabine Markowski and Lisa Zeller (both Institute of Clinical Microbiology and Hygiene, Regensburg, University Hospital Regensburg, Germany), Claudia Fischer (Institute of Pathology, University of Regensburg, Germany) as well as Michelle Wröbel and Ursula Schombel (both Research Center Borstel, Germany) for technical support. We thank Dr. Uwe Mamat (Research Center Borstel, Germany) for providing the *Pa* PAO1 strain.

## Supplementary Figures

**Figure S1:**
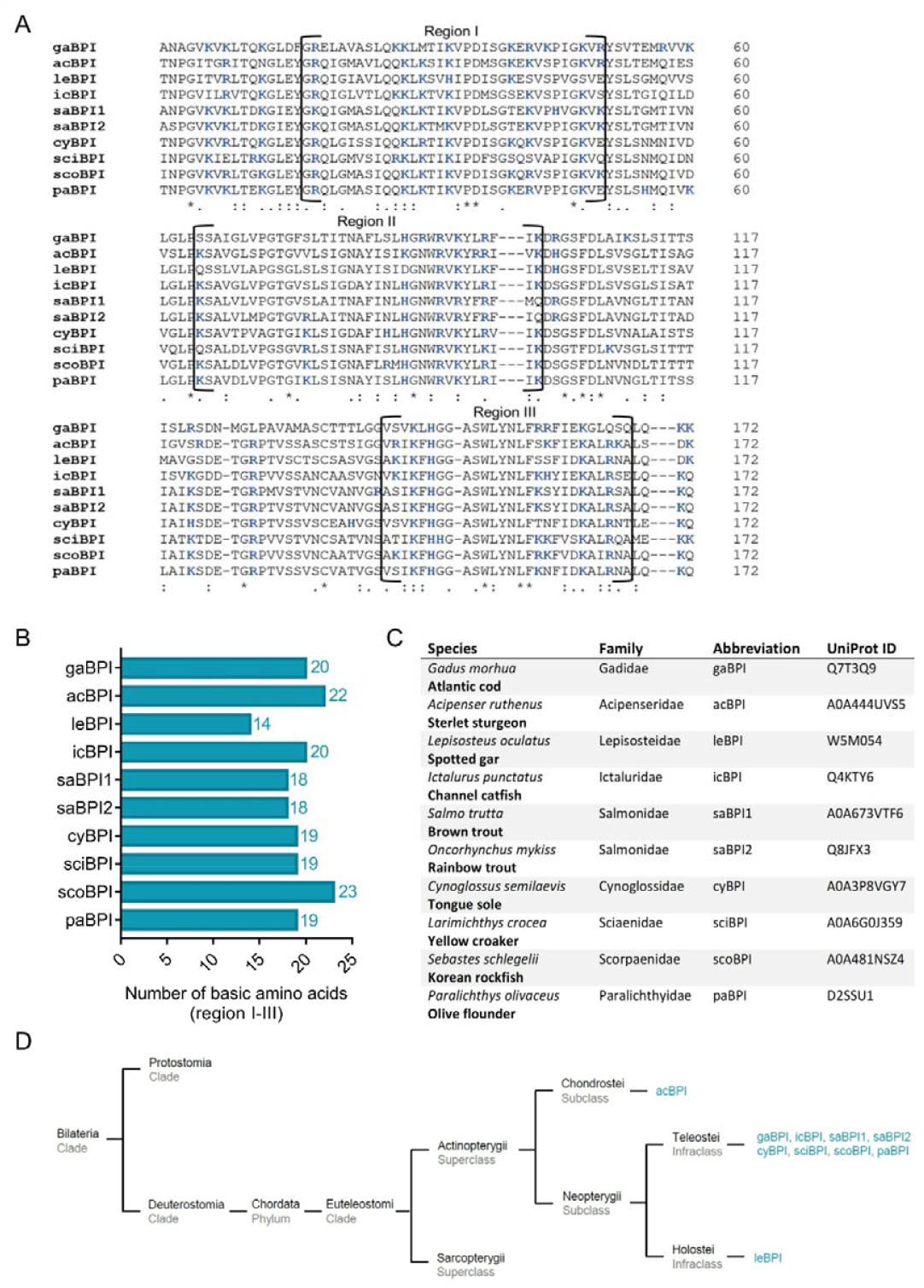
Sequence alignment and analysis of orthologous Actinopterygii BPI. (**A**) Sequence alignment of amino acid sequences of orthologous proteins. Functional regions I, II and III are framed and positively charged amino acids are shown in blue. Conserved areas are indicated by asterisk, colon and period representing highly, moderately and less conserved areas, respectively. (**B**) Number of basic amino acids in regions I–III of Actinopterygii BPI. (**C**) Actinopterygii species are listed including taxonomic familiy, abbreviation of the corresponding BPI and UniProt ID. (**D**) Phylogenetic tree analysis mapping Actinopterygii BPI orthologous to their respective taxonomic ranks.

**Figure S2:**
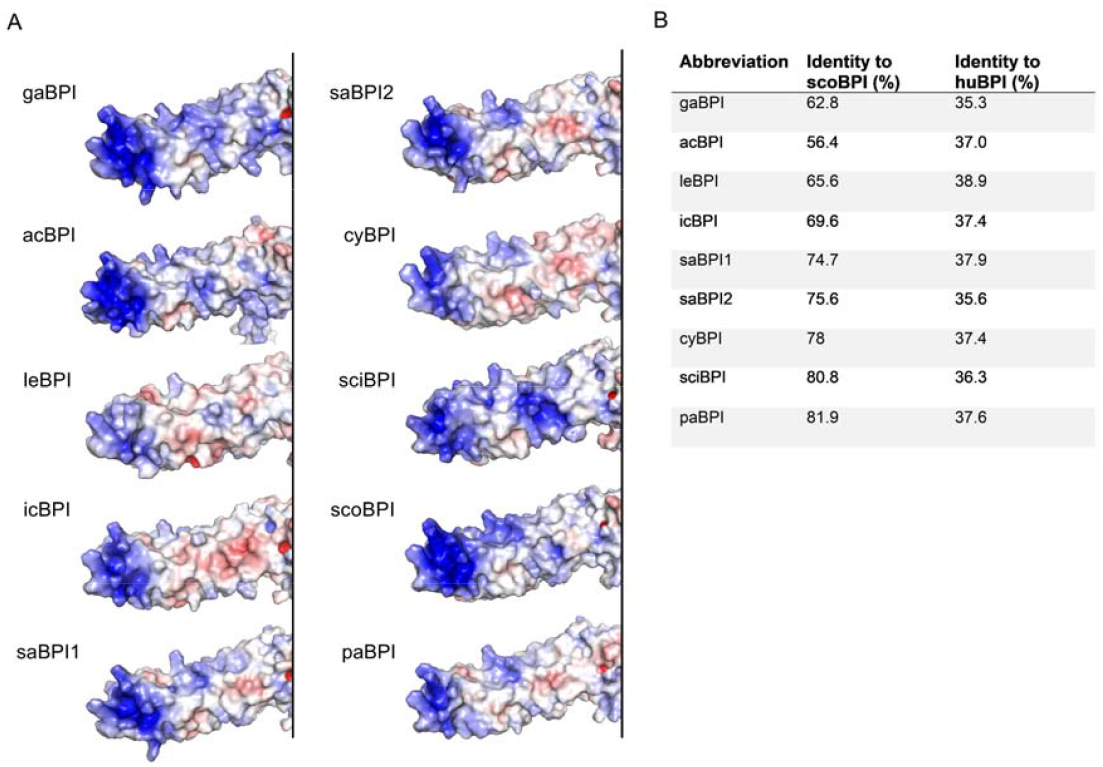
Predicted N-terminal surface electrostatics and sequence identities of Actinopterygii BPI. (**A**) Surface electrostatics for the N-terminal barrels of three-dimensional structure predictions for BPI of investigated Actinopterygii. Areas colored in blue represent positively charged areals an, negatively charged domains are shown in red. (**B**) Sequence identities of investigated orthologous proteins to scoBPI and huBPI.

**Figure S3:**
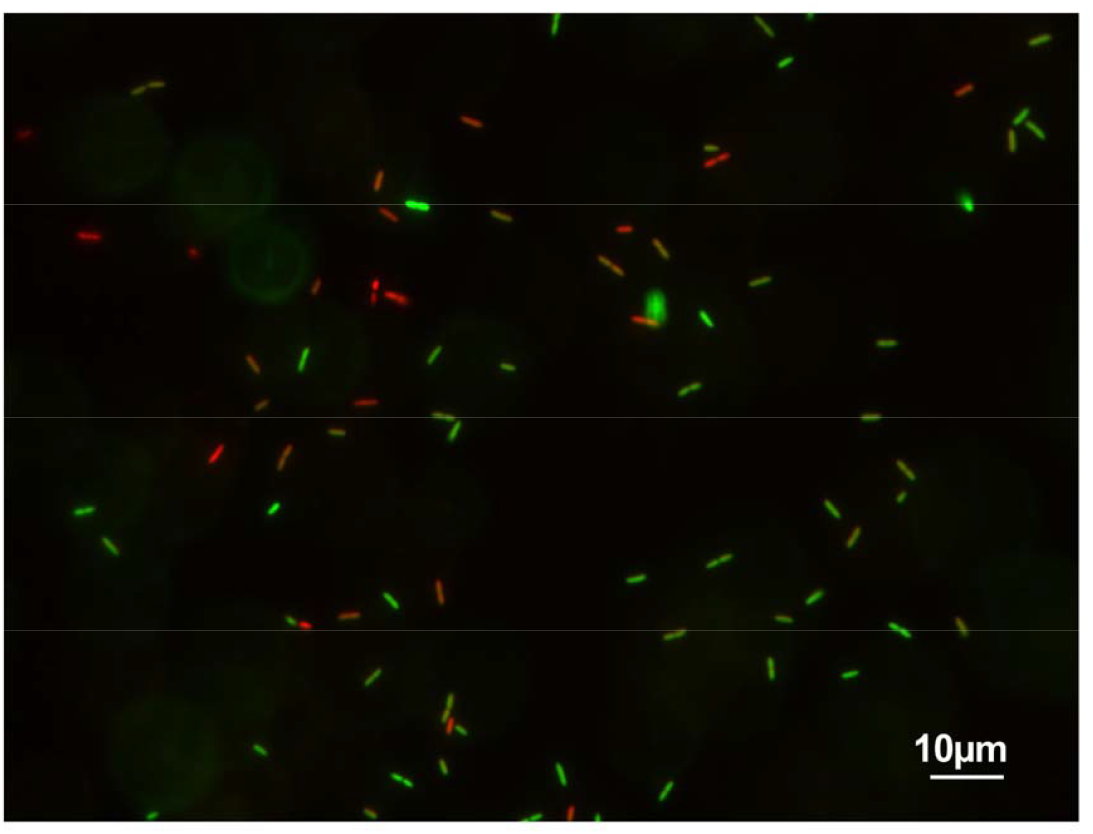
Corresponding microscopic imaging of viability in *Pa* MDR at the time of TEM imaging. Representative image of bacterial viability assay performed with scoBPI-treated *Pa* MDR strain at the time of TEM. Viable bacteria are seen in green and dead bacteria in red, respectively. Bar scales for 10 μm.

## References

1. Collins FS. Cystic fibrosis: Molecular Biology and Therapeutic Implications. Science. 1992;256(5058):774–779. doi: 10.1126/science.256.5058.774.

2. Boucher RC. Airway surface dehydration in cystic fibrosis: pathogenesis and therapy. Annu Rev Med. 2007;58:157–170. doi: 10.1146/annurev.med.58.071905.105316.

3. Gustafsson JK, et al. Bicarbonate and functional CFTR channel are required for proper mucin secretion and link cystic fibrosis with its mucus phenotype. J Exp Med. 2012;209(7):1263–1272. doi: 10.1084/jem.20120562.

4. Middleton PG, et al. Elexacaftor-Tezacaftor-Ivacaftor for Cystic Fibrosis with a Single Phe508del Allele. N Engl J Med. 2019;381(19):1809–1819. doi: 10.1056/NEJMoa1908639.

5. Koch C. Early infection and progression of cystic fibrosis lung disease. Pediatr Pulmonol. 2002;34(3):232–236. doi: 10.1002/ppul.10135.

6. Emerson J, Rosenfeld M, McNamara S, Ramsey B, Gibson RL. *Pseudomonas aeruginosa* and other predictors of mortality and morbidity in young children with cystic fibrosis. Pediatr Pulmonol. 2002;34(2):91–100. doi: 10.1002/ppul.10127.

7. Balaban NQ, et al. Definitions and guidelines for research on antibiotic persistence. Nat Rev Microbiol. 2019;17(7):441–448. doi: 10.1038/s41579-019-0196-3.

8. Rossi E, et al. *Pseudomonas aeruginosa* adaptation and evolution in patients with cystic fibrosis. Nat Rev Microbiol. 2021;19(5):331–342. doi: 10.1038/s41579-020-00477-5.

9. Pitt TL, Sparrow M, Warner M, Stefanidou M. Survey of resistance of *Pseudomonas aeruginosa* from UK patients with cystic fibrosis to six commonly prescribed antimicrobial agents. Thorax. 2003;58(9):794–796. doi: 10.1136/thorax.58.9.794.

10. Weiss J, Elsbach P, Olsson I, Odeberg H. Purification and characterization of a potent bactericidal and membrane active protein from the granules of human polymorphonuclear leukocytes. J Biol Chem. 1978;253(8):2664–2672.

11. Weiss J, et al. Human bactericidal/permeability-increasing protein and a recombinant NH2-terminal fragment cause killing of serum-resistant gram-negative bacteria in whole blood and inhibit tumor necrosis factor release induced by the bacteria. J Clin Invest. 1992;90(3):1122–1130. doi: 10.1172/JCI115930.

12. Dentener MA, Asmuth EJ von, Francot GJ, Marra MN, Buurman WA. Antagonistic effects of lipopolysaccharide binding protein and bactericidal/permeability-increasing protein on lipopolysaccharide-induced cytokine release by mononuclear phagocytes. Competition for binding to lipopolysaccharide. J Immunol. 1993;151(8):4258–4265.

13. Weiss J, Victor M, Elsbach P. Role of charge and hydrophobic interactions in the action of the bactericidal/permeability-increasing protein of neutrophils on gram-negative bacteria. J Clin Invest. 1983;71(3):540–549. doi: 10.1172/jci110798.

14. Ederer KU, et al. A Polymorphism of Bactericidal/Permeability-Increasing Protein Affects Its Neutralization Efficiency towards Lipopolysaccharide. Int J Mol Sci. 2022;23(3). doi: 10.3390/ijms23031324.

15. Weitz A, Spotnitz R, Collins J, Ovadia S, Iovine NM. Log reduction of multidrug-resistant Gram-negative bacteria by the neutrophil-derived recombinant bactericidal/permeability-increasing protein. Int J Antimicrob Agents. 2013;42(6):571–574. doi: 10.1016/j.ijantimicag.2013.07.019.

16. Theprungsirikul J, Skopelja-Gardner S, Rigby WFC. Killing three birds with one BPI: Bactericidal, opsonic, and anti-inflammatory functions. J Transl Autoimmun. 2021;4:100105. doi: 10.1016/j.jtauto.2021.100105.

17. Schultz H, Schinke S, Mosler K, Herlyn K, Schuster A, Gross WL. BPI-ANCA of pediatric cystic fibrosis patients can impair BPI-mediated killing of *E. coli* DH5alpha in vitro. Pediatr Pulmonol. 2004;37(2):158–164. doi: 10.1002/ppul.10416.

18. McQuillan K, Gargoum F, Murphy MP, McElvaney OJ, McElvaney NG, Reeves EP. Targeting IgG Autoantibodies for Improved Cytotoxicity of Bactericidal Permeability Increasing Protein in Cystic Fibrosis. Front. Pharmacol. 2020;11:1098. doi: 10.3389/fphar.2020.01098.

19. Hovold G, Lindberg U, Ljungberg JK, Shannon O, Påhlman LI. BPI-ANCA is expressed in the airways of cystic fibrosis patients and correlates to platelet numbers and *Pseudomonas aeruginosa* colonization. Respir Med. 2020;170:105994. doi: 10.1016/j.rmed.2020.105994.

20. Carlsson M, et al. Autoantibody response to BPI predict disease severity and outcome in cystic fibrosis. J Cyst Fibr. 2007;6(3):228–233. doi: 10.1016/j.jcf.2006.10.005.

21. Wittmann I, Schönefeld M, Aichele D, Groer G, Gessner A, Schnare M. Murine bactericidal/permeability-increasing protein inhibits the endotoxic activity of lipopolysaccharide and gram-negative bacteria. J Immunol. 2008;180(11):7546–7552. doi: 10.4049/jimmunol.180.11.7546.

22. Lee S, Elvitigala DAS, Lee S, Kim HC, Park H-C, Lee J. Molecular characterization of a bactericidal permeability-increasing protein/lipopolysaccharide-binding protein from black rockfish (Sebastes schlegelii): Deciphering its putative antibacterial role. Dev Comp Immunol. 2017;67:266–275. doi: 10.1016/j.dci.2016.09.011.

23. Gonzalez M, et al. Evidence of a bactericidal permeability increasing protein in an invertebrate, the Crassostrea gigas Cg-BPI. PNAS. 2007;104(45):17759–17764. doi: 10.1073/pnas.0702281104.

24. Zhang Y, He X, Li X, Fu D, Chen J, Yu Z. The second bactericidal permeability increasing protein (BPI) and its revelation of the gene duplication in the Pacific oyster, Crassostrea gigas. Fish Shellfish Immunol. 2011;30(3):954–963. doi: 10.1016/j.fsi.2011.01.031.

25. Little RG, Kelner DN, Lim E, Burke DJ, Conlon PJ. Functional domains of recombinant bactericidal/permeability increasing protein (rBPI23). J Biol Chem. 1994;269(3):1865–1872.

26. Gazzano-Santoro H, et al. Characterization of the structural elements in lipid A required for binding of a recombinant fragment of bactericidal/permeability-increasing protein rBPI23. Infect Immun. 1995;63(6):2201–2205. doi: 10.1128/iai.63.6.2201-2205.1995.

27. Sun Y-Y, Sun L. A Teleost Bactericidal Permeability-Increasing Protein Kills Gram-Negative Bacteria, Modulates Innate Immune Response, and Enhances Resistance against Bacterial and Viral Infection. PloS one. 2016;11(4):e0154045. doi: 10.1371/journal.pone.0154045.

28. Beamer LJ, Carroll SF, Eisenberg D. Crystal structure of human BPI and two bound phospholipids at 2.4 angstrom resolution. Science. 1997;276(5320):1861–1864. doi: 10.1126/science.276.5320.1861.

29. Jumper J, et al. Highly accurate protein structure prediction with AlphaFold. Nature. 2021;596(7873):583–589. doi: 10.1038/s41586-021-03819-2.

30. Varadi M, et al. AlphaFold Protein Structure Database: massively expanding the structural coverage of protein-sequence space with high-accuracy models. Nucleic Acids Res. 2022;50(D1):D439–D444. doi: 10.1093/nar/gkab1061.

31. Aichele D, Schnare M, Saake M, Röllinghoff M, Gessner A. Expression and antimicrobial function of bactericidal permeability-increasing protein in cystic fibrosis patients. Infect Immun. 2006;74(8):4708–4714. doi: 10.1128/IAI.02066-05.

32. Bülow S, et al. Lipopolysaccharide Binding Protein and Bactericidal/Permeability-Increasing Protein as Biomarkers for Invasive Pulmonary Aspergillosis. J Fungi. 2020;6(4):304. doi: 10.3390/jof6040304.

33. Tobias PS, Soldau K, Iovine NM, Elsbach P, Weiss J. Lipopolysaccharide (LPS)-binding proteins BPI and LBP form different types of complexes with LPS. J Biol Chem. 1997;272(30):18682–18685. doi: 10.1074/jbc.272.30.18682.

34. Wiese A, Brandenburg K, Carroll SF, Rietschel ET, Seydel U. Mechanisms of action of bactericidal/permeability-increasing protein BPI on reconstituted outer membranes of gram-negative bacteria. Biochemistry. 1997;36(33):10311–10319. doi: 10.1021/bi970177e.

35. Mojsoska B, et al. The high persister phenotype of *Pseudomonas aeruginosa* is associated with increased fitness and persistence in cystic fibrosis airways. 2019. doi: 10.1101/561589.

36. Łapińska U, et al. Fast bacterial growth reduces antibiotic accumulation and efficacy. eLife. 2022;11. doi: 10.7554/eLife.74062.

37. Malhotra S, Limoli DH, English AE, Parsek MR, Wozniak DJ. Mixed Communities of Mucoid and Nonmucoid *Pseudomonas aeruginosa* Exhibit Enhanced Resistance to Host Antimicrobials. mBio. 2018;9(2). doi: 10.1128/mBio.00275-18.

38. Cabral DA, Loh BA, Speert DP. Mucoid *Pseudomonas aeruginosa* resists nonopsonic phagocytosis by human neutrophils and macrophages. Pediatr Res. 1987;22(4):429–431. doi: 10.1203/00006450-198710000-00013.

39. Høiby N, Ciofu O, Bjarnsholt T. *Pseudomonas aeruginosa* biofilms in cystic fibrosis. Future Microbiol. 2010;5(11):1663–1674. doi: 10.2217/fmb.10.125.

40. Rice LB. Federal funding for the study of antimicrobial resistance in nosocomial pathogens: no ESKAPE. J Infect Dis. 2008;197(8):1079–1081. doi: 10.1086/533452.

41. Zhao MH, Jones SJ, Lockwood CM. Bactericidal/permeability-increasing protein (BPI) is an important antigen for anti-neutrophil cytoplasmic autoantibodies (ANCA) in vasculitis. Clin Exp Immunol. 1995;99(1):49–56. doi: 10.1111/j.1365-2249.1995.tb03471.x.

42. Manolova I, Dancheva M, Halacheva K. Antineutrophil cytoplasmic antibodies in patients with systemic lupus erythematosus: prevalence, antigen specificity, and clinical associations. Rheumatol Int. 2001;20(5):197–204. doi: 10.1007/s002960100108.

43. Teh CL, Wan SA, Ling GR. Severe infections in systemic lupus erythematosus: disease pattern and predictors of infection-related mortality. Clin Rheumatol. 2018;37(8):2081–2086. doi: 10.1007/s10067-018-4102-6.

44. Levin M, et al. Recombinant bactericidal/permeability-increasing protein (rBPI21) as adjunctive treatment for children with severe meningococcal sepsis: a randomised trial. The Lancet. 2000;356(9234):961–967. doi: 10.1016/S0140-6736(00)02712-4.

45. Möhlen MA von der, et al. Inhibition of endotoxin-induced cytokine release and neutrophil activation in humans by use of recombinant bactericidal/permeability-increasing protein. J Infect Dis. 1995;172(1):144–151. doi: 10.1093/infdis/172.1.144.

46. Cantin AM, Hartl D, Konstan MW, Chmiel JF. Inflammation in cystic fibrosis lung disease: Pathogenesis and therapy. J Cyst Fibr. 2015;14(4):419–430. doi: 10.1016/j.jcf.2015.03.003.

47. Kelly E, Greene CM, McElvaney NG. Targeting neutrophil elastase in cystic fibrosis. Expert Opin Ther Targets. 2008;12(2):145–157. doi: 10.1517/14728222.12.2.145.

48. Taggart C, Coakley RJ, Greally P, Canny G, O’Neill SJ, McElvaney NG. Increased elastase release by CF neutrophils is mediated by tumor necrosis factor-alpha and interleukin-8. Am J Physiol Lung Cell Mol Physiol. 2000;278(1):L33–41. doi: 10.1152/ajplung.2000.278.1.L33.

49. Zhang P-X, et al. Reduced caveolin-1 promotes hyperinflammation due to abnormal heme oxygenase-1 localization in lipopolysaccharide-challenged macrophages with dysfunctional cystic fibrosis transmembrane conductance regulator. J Immunol. 2013;190(10):5196–5206. doi: 10.4049/jimmunol.1201607.

50. Iovine NM, Elsbach P, Weiss J. An opsonic function of the neutrophil bactericidal/permeability-increasing protein depends on both its N- and C-terminal domains. PNAS. 1997;94(20):10973–10978. doi: 10.1073/pnas.94.20.10973.

51. Theprungsirikul J, Skopelja-Gardner S, Burns AS, Wierzbicki RM, Rigby WFC. Bactericidal/Permeability-Increasing Protein Preeminently Mediates Clearance of *Pseudomonas aeruginosa* In Vivo via CD18-Dependent Phagocytosis. Front. Immunol. 2021;12:659523. doi: 10.3389/fimmu.2021.659523.

52. Bülow S, et al. Bactericidal/Permeability-Increasing Protein Is an Enhancer of Bacterial Lipoprotein Recognition. Front Immunol. 2018;9:2768. doi: 10.3389/fimmu.2018.02768.

53. Kutschera A, et al. Bacterial medium-chain 3-hydroxy fatty acid metabolites trigger immunity in Arabidopsis plants. Science. 2019;364(6436):178–181. doi: 10.1126/science.aau1279.

54. Westphal O, Jann K. Bacterial Lipopolysaccharides Extraction with Phenol-Water and Further Applications of the Procedure. Methods in Carbohydrate Chemistry: General polysaccharides:83–91.

55. Jimenez-Barbero J, Castro Cd, Evidente A, Molinaro A, Parrilli M, Surico G. Structural Determination of the O-Specific Chain of the Lipopolysaccharide from *Pseudomonas cichorii*. Eur J Org Chem. 2002;2002(11):1770–1775. doi: 10.1002/1099-0690(200206)2002:11<1770:AID-EJOC1770>3.0.CO;2-5.

56. The UniProt Consortium, et al. UniProt: the universal protein knowledgebase in 2021. Nucleic Acids Res. 2021;49(D1):D480–D489. doi: 10.1093/nar/gkaa1100.

57. Sievers F, et al. Fast, scalable generation of high-quality protein multiple sequence alignments using Clustal Omega. Mol Syst Biol. 2011;7:539. doi: 10.1038/msb.2011.75.

58. Huang X, Miller W. A time-efficient, linear-space local similarity algorithm. Adv Appl Math. 1991;12(3):337–357. doi: 10.1016/0196-8858(91)90017-D.

